# Variability in gene expression is associated with incomplete penetrance in inherited eye disorders

**DOI:** 10.1101/2020.01.28.915504

**Authors:** David J. Green, Shalaw R. Sallah, Jamie M. Ellingford, Simon C. Lovell, Panagiotis I. Sergouniotis

## Abstract

Inherited eye disorders (IED) are a heterogeneous group of Mendelian conditions that are associated with visual impairment. Although these disorders often exhibit incomplete penetrance and variable expressivity, the scale and mechanisms of these phenomena remain largely unknown. Here, we utilize publicly-available genomic and transcriptomic datasets to gain insights into variable penetrance in IED. Variants in a curated set of 340 IED-implicated genes were extracted from HGMD 2019.1 and cross-checked with the gnomAD 2.1 control-only dataset. Genes for which >1 variant was encountered in both HGMD and gnomAD were considered to be associated with variable penetrance (n=56). Variability in gene expression levels was then estimated for the subset of these genes that was found to be adequately expressed in two relevant resources, GTEx and EyeGEx. We found that genes suspected to be associated with variable penetrance tended to have significantly more variability in gene expression levels in the general population (p=0.0000015); this finding was consistent across tissue types. The results of this study point to a possible influence of *cis* and/or *trans*-acting elements on the expressivity of variants causing Mendelian disorders. They also highlight the potential utility of quantifying gene expression as part of the investigation of families showing evidence of variable penetrance.

## Introduction

Incomplete penetrance and variable expressivity (hereafter referred to as variable penetrance) are two closely linked biological phenomena leading to non-linearity between genotype and phenotype. Incomplete penetrance occurs when individuals with a particular disease-predisposing genetic change fail to express the corresponding disease phenotype; variable expressivity occurs when individuals carrying identical genetic changes display significant variability in terms of disease severity and/or onset. These phenomena are widespread in human genetics, even in the context of Mendelian disorders, *i.e.* conditions that are driven by monoallelic or biallelic variants with strong effects [1]. Importantly, a firm understanding of variable penetrance is required to improve the predictive power of genomic data and to enable accurate variant interpretation [2,3].

The mechanisms underlying variable penetrance remain largely unknown [1]. While multiple genetic and environmental factors may contribute, variability in gene expression is increasingly recognized as an important source of variable phenotypic expression [4–8]. Emerging resources combining large-scale genomic and transcriptomic datasets are making analysis of these phenomena increasingly tractable. For example, a recent study using data from the Genotype-Tissue Expression Project (GTEx) [9] has shown that, in the general population, purifying selection has depleted haplotype combinations predicted to increase the penetrance of likely disease-associated variants (*i.e*. combinations of a regulatory variant associated with higher haplotype expression and a variant predicted to be pathogenic by *in silico* tools) [6]. Another study utilising genomic and transcriptomic data from healthy retinal donors found evidence for *cis*-acting expression regulation in genes that are implicated in retinal disorders exhibiting variable penetrance [10]. Notably, both these studies focused on *cis*-acting effects despite the fact that these account only for a fraction of gene expression variability [11].

Here, we integrate publicly available genomic and transcriptomic datasets to study variable penetrance in a well-characterized group of Mendelian conditions, inherited eye disorders (IED). The central hypothesis driving this work is that IED-implicated genes associated with variable phenotypic expression in patients will exhibit variable gene expression in the general population. To test this hypothesis, we: [i] identified a subset of IED-implicated genes that are enriched for variants with possible variable penetrance (IP/VE-associated genes) and [ii] compared gene expression levels between these genes and other IED-implicated genes for which there was no evidence of variable penetrance.

## Materials and Methods

### Obtaining lists of genes and variants associated with inherited eye disorders

IED-implicated genes were obtained through PanelApp, a publicly available knowledgebase that combines expert reviews and manual curation to create diagnostic-grade gene lists [12]. We focused on the genes that are included in the different panels within the “Ophthalmological disorders” disease category (as of July 2019): [i] anophthalmia or microphthalmia panel (version 1.21), [ii] cataract panel (version 1.27), [iii] corneal abnormalities panel (version 1.7), [iv] developmental glaucoma panel (version 1.5), [v] infantile nystagmus panel (version 1.3), [vi] ocular coloboma panel (version 1.34), [vii] optic neuropathy panel (version 1.116) and [viii] retinal disorders panel (version 1.145). Only genes with a high level of evidence (“green”) were retained, as our aim was to study variable penetrance in genes robustly associated with disease (rather than to question gene-disease associations). Information on the mode of inheritance relevant to each gene was also obtained from PanelApp; the following categories were included: monoallelic (corresponding to autosomal dominant), biallelic (corresponding to autosomal recessive), both monoallelic and biallelic (corresponding to both autosomal dominant and autosomal recessive) and X-linked (corresponding to X-linked dominant or X-linked recessive).

Disease-associated variants in the above IED-implicated genes were obtained from the Human Gene Mutation Database (HGMD) Professional (version 2019.1, accessed June 2019). HGMD represents an attempt to collate all published variants that underlie, or are closely associated with human inherited disease [13]. At present, it is the largest resource of its kind including >250,000 disease-associated variants. Only variants labelled as “disease mutation” (DM, indicating the highest level of clinical significance) were retained. The following variant types were selected: missense, nonsense, splicing, small indel, small deletion, and small insertion. The build 37 (GRCh37) genomic coordinates of all variants were obtained using Variant Validator version 0.1.3 [14]. At this point, we excluded genes with only one entry with the DM flag in HGMD 2019.1 (n=8; *RP9*, *TUB*, *TMEM126A*, *IDH3B*, *FOXD3*, *SSBP1*, *MIR184*, *OVOL2*).

To limit the potential impact of annotation errors in HGMD, we obtained the CADD PHRED-scaled score (a widely used measure of variant deleteriousness) for each variant, and only considered changes with a score >15 as disease-associated. CADD is an integrative annotation tool built from more than 60 genomic features. A PHRED-scaled score ≥10 indicates a raw score in the top 10% of all possible single nucleotide variants, while a score of ≥20 indicates a raw score in the top 1% [15]. Notably, the CADD team proposed (but did not recommend for categorical usage) 15 as a possible cut-off value, although they highlighted the arbitrary nature of such a cut-off [16]. CADD version 1.4 (GRCh37) was used in this study (accessed July 2019).

### Identifying genes that are possibly associated with variable penetrance

Initially, we aimed to identify IED-associated variants that are present in unaffected individuals. To achieve this, we looked for HGMD-listed changes that have a CADD score >15 and are present in the Genome Aggregation Database (gnomAD). gnomAD incorporates exome or genome sequence data from >140,000 humans; these data were obtained primarily from case-control studies of adult-onset common diseases, including cardiovascular disease, type 2 diabetes, and psychiatric disorders [17]. The gnomAD 2.1 release was used (accessed June 2019) and we focused on the control-only subset, which includes samples from >60,000 individuals who were not selected as a case in a case/control study of common disease. It is noteworthy that, by design, gnomAD aims to exclude individuals with severe paediatric diseases from the released dataset [17]. Cross-checking between HGMD 2019.1 and gnomAD 2.1 was performed based on the genomic co-ordinates, and reference and alternate alleles for each variant.

Subsequently, we aimed to identify a subset of IED-implicated genes that are possibly associated with incomplete penetrance and/or variable expressivity (IP/VE-associated genes). The following criteria were used to define IP/VE-associated genes: [i] genes classified as green and monoallelic in PanelApp for which at least two HGMD-listed variants (with a CADD score >15) were found to be present in heterozygous state in at least two individuals from the gnomAD control-only dataset; [ii] genes classified as green and biallelic in PanelApp for which at least two HGMD-listed variants (with a CADD score >15) were found to be present in homozygous state in at least two individuals from the gnomAD control-only dataset; [iii] genes classified as green and both monoallelic and biallelic in PanelApp for which at least two HGMD-listed variants (with a CADD score >15) were found to be present in homozygous state (in the case of variants reported to cause disease through a recessive mode of inheritance after manual inspection of original reports) or heterozygous state (in the case of variants reported to cause disease through a dominant mode of inheritance after manual inspection of original reports) in at least two individuals from the gnomAD control-only dataset; [iv] genes classified as green and X-linked in PanelApp for which at least two HGMD-listed variants (with a CADD score >15) were found to be present in either hemizygous or homozygous state in at least two individuals from the gnomAD control-only dataset.

We used a cut-off of two individuals to account for [i] contamination of the gnomAD control dataset with individuals who have Mendelian disorders, [ii] sequencing errors, and [iii] the presence of a disease-predisposing variant in a gnomAD control individual who is too young to be symptomatic. A flow-chart outlining the classification of genes into IP/VE-associated and other is shown in Figure 1. Notably, secondary analyses were performed using the same approach but adopting more stringent variant-level cut-offs, including a CADD score >20 and a CADD score >20 plus a minor allele frequency of <0.01 (in gnomAD 2.1).

**Figure 1.**
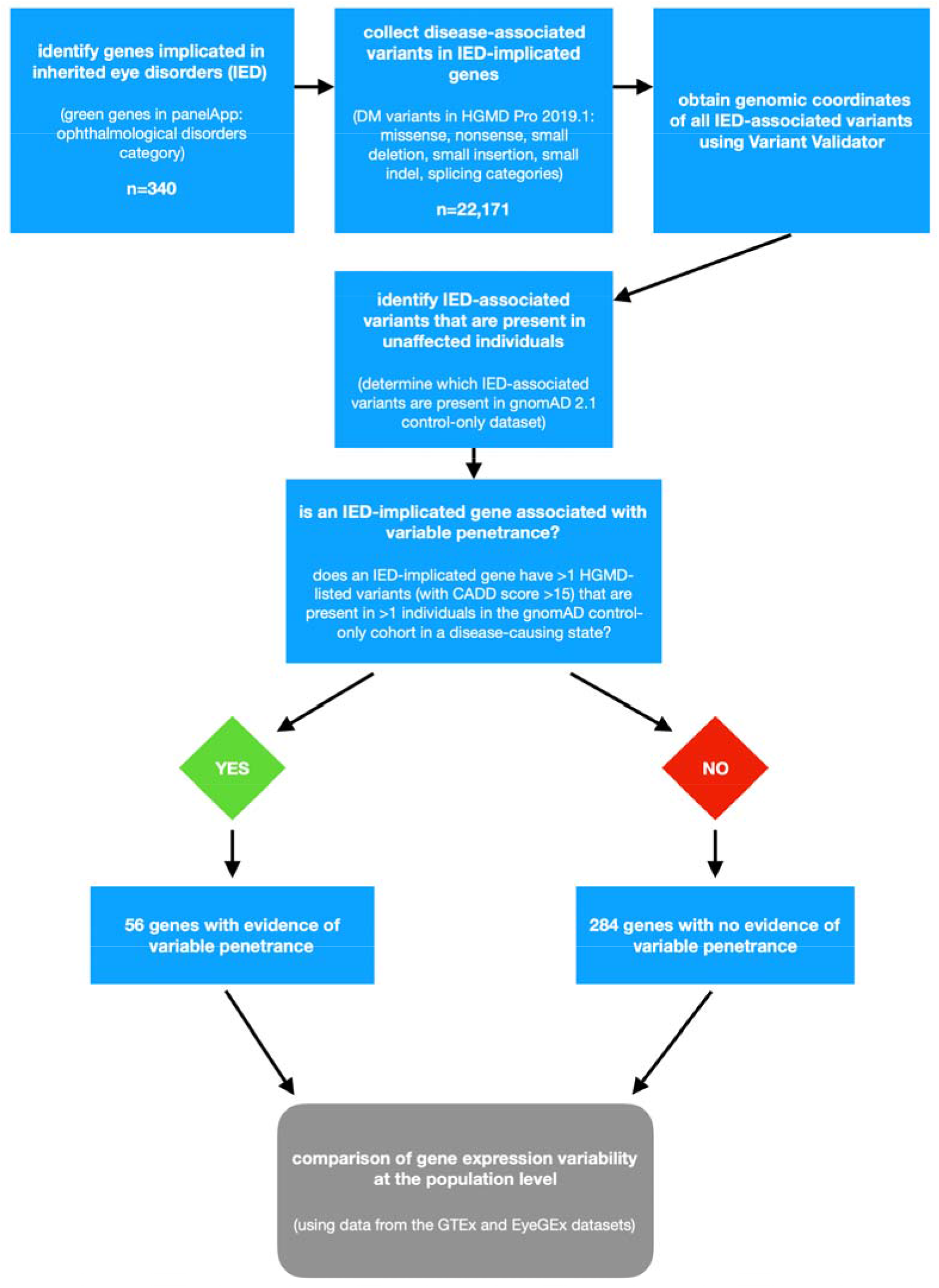
Flowchart outlining the study design of the project. IED, inherited eye disorders; HGMD, Human Gene Mutation Database; gnomAD, Genome Aggregation Database; CADD, Combined Annotation-Dependent Depletion; GTEx, Genotype-Tissue Expression Project; EyeGEx, Eye Genotype Expression.

Two gene expression resources were used to determine variability in population mRNA levels for IED-implicated genes: the GTEx [9] and the Eye Genotype Expression (EyeGEx) [18] databases. The GTEx v8 dataset (accessed September 2019) consisting of 948 donors and 17,382 samples from 52 tissues and two cell lines was used in this study. Given that most IED-implicated genes are linked to retinal disease (187/340) and that the GTEx v8 release does not include retinal tissue, we chose to also analyse data from the EyeGEx dataset. EyeGEx includes RNA-sequencing data from 453 post-mortem peripheral retina samples (105 control samples and 348 samples from donors with age-related macular degeneration) all processed in a uniform manner [18].

Variability in mRNA levels in the population was assessed for the entire GTEx v8 dataset using a method similar to that developed by Simonovsky *et al.* and using parameters broadly in keeping with those used by the GTEx Consortium [19]. First, raw gene read data were downloaded from the GTEx web server and normalized to obtain the same library size for every sample for each tissue using the trimmed mean of M-values (TMM) method implemented in edgeR [20]. Prior to normalization, genes with at most 10 reads in all samples were removed to exclude lowly expressed genes, as these were considered to represent noise [19]. A comprehensive list of protein-coding genes was obtained from Ensembl Biomart [21]. The local coefficient of variation (LCV), a measure that is strongly correlated with other measures of gene expression variability (including expression variation [22] and distance from the median coefficient of variation [23]) but more robust to lacking data and not correlated with gene expression level, was calculated for all genes. To ensure that genes were adequately expressed in enough samples to allow the estimation of gene expression variability, we focused on genes expressed to at least 7 cpm in at least 80% of samples in a tissue for all tissues with at least 20 samples. This resulted in variability estimates for 15,398 genes across 49 tissues in the entire GTEx dataset based on a total of 17,176 samples. Median variability across all available tissues (including brain tissues and skin tissues) was calculated.

The same procedure was performed for the single-tissue EyeGEx dataset. Raw estimated gene reads (output from RMEM) were downloaded from the GTEx web server and normalized in the same manner as the GTEx data, as described above. The LCV was calculated for all genes expressed at 7 cpm in at least 80% of retinal samples. This yielded variability estimates for 9,942 genes based on a total of 453 samples. Tables containing the calculated LCV scores for all genes across the GTEx and EyeGEx datasets are provided in Supplementary File 1. The correlation between the EyeGEx LCV score and the GTEx (median across all tissues) LCV score was evaluated in the subset of genes that were found to be adequately expressed in both datasets (Supplementary File 2).

Variability in mRNA levels was subsequently compared between IP/VE-associated genes and all other IED-implicated genes. Genes implicated in retinal disorders were also assessed separately using data from the EyeGEx resource. All statistical analyses were performed using Python 3 (https://www.python.org) and p-values were computed conventionally (findings reported in the text and Table 1) and using resampling without replacement (findings reported in Table 1). The latter involved randomly reshuffling the group labels (for each group listed in Table 1) 1,000,000 times and calculating how often the observed difference occurred using the permutation_test function in the Python package mlxtend. The presented permutation p-values represent the number of positive findings divided by the total number of permutations; thus, a p-value of 0.05 indicates that the observed difference occurred in 5% of permutations.

**Table 1.** Comparison of gene expression variability between genes that are possibly associated with variable penetrance and genes for which no evidence of variable penetrance was detected.

| panelApp gene set | expression dataset used | LCV for IP/VE IED genes<sup>(number of genes)</sup> | LCV for other IED genes<sup>(number of genes)</sup> | p-value | permutation p-value |
| --- | --- | --- | --- | --- | --- |
| all genes implicated in ophthalmological disorders | GTEx all tissues | 83.7<sup>(43)</sup> | 48.85<sup>(216)</sup> | 0.0000015 | 0.00002 |
| all genes implicated in ophthalmological disorders | GTEx brain tissues | 80.4<sup>(35)</sup> | 49.25<sup>(184)</sup> | 0.0000053 | 0.0001 |
| all genes implicated in ophthalmological disorders | GTEx whole blood | 66.2<sup>(15)</sup> | 40.2<sup>(101)</sup> | 0.018 | 0.044 |
| all genes implicated in ophthalmological disorders | GTEx sun-exposed skin | 79.9<sup>(22)</sup> | 38.9<sup>(156)</sup> | 0.0000083 | 0.00001 |
| all genes implicated in ophthalmological disorders | GTEx non-sun-exposed skin | 82<sup>(23)</sup> | 36.9<sup>(152)</sup> | 0.000017 | 0.000002 |
| all genes implicated in ophthalmological disorders | GTEx cultured fibroblasts | 78.4<sup>(23)</sup> | 43.8<sup>(152)</sup> | 0.00001 | 0.0004 |
| genes implicated in retinal disorders only | EyeGEx | 83.8<sup>(23)</sup> | 67.8<sup>(137)</sup> | 0.0028 | 0.0021 |

IED, inherited eye disorders; LCV for IP/VE IED genes, median local coefficient of variation for IED-implicated genes with evidence for incomplete penetrance and/or variable expressivity; LCV for other IED genes, median local coefficient of variation for IED-implicated genes for which no evidence of incomplete penetrance was found. GTEx, Genotype-Tissue Expression Project version 8 dataset [9]; EyeGEx, Eye Genotype Expression (EyeGEx) dataset [18]; permutation p-value, the result of permutation analysis (please see the Methods section for further information).

A potential confounder in these analyses is coding constraint, *i.e*. the strength of natural selection acting on each gene/region [24]. The following constraint metrics were obtained from gnomAD 2.1 for each studied gene: missense z-score [25], missense observed/expected (o/e) ratio, loss-of-function o/e ratio and loss-of-function o/e upper bound fraction (LOEUF) [17]. The relationship between these constraint metrics and gene expression variability (GTEx and EyeGEx LCV) was subsequently studied. The difference in constraint between different groups of genes (IP/VE-associated genes and all other IED-implicated genes) was also investigated (Supplementary File 2).

## Results

### Variants associated with inherited eye disorders are frequently encountered in unaffected individuals

Overall, 340 genes with a high level of evidence for disease-causation were found in the “Ophthalmological disorders” category of PanelApp. This included 183 genes associated with inherited retinal disorders, 89 genes associated with cataract and 85 genes associated with other ophthalmic conditions; 9% of genes (32/340) were linked to more than one disorder (see Supplementary File 3 for more information). In terms of mode of inheritance, 81 genes were labelled as monoallelic, 201 genes were labelled as biallelic, 41 genes were labelled as both monoallelic and biallelic, and 17 genes were labelled as X-linked.

A total of 22,171 variants across the 340 IED-implicated genes were identified in HGMD 2019.1 (missense, nonsense, splicing, small indel, small deletion, and small insertion categories); 46% (10,188/22,171) were missense and 17% (3,755/22,171) were nonsense changes. In terms of mode of inheritance (according to PanelApp), 24% (5,342/22,171) of variants were in genes labelled as monoallelic, 51% (11,316/22,171) in genes labelled as biallelic, 16% (3,437/22,171) in genes labelled as both monoallelic and biallelic, and 9% (2,076/22,171) in genes labelled as X-linked.

In total, 1.3% (285/22,171) of these IED-associated variants were present in at least two individuals in the gnomAD control-only cohort in a disease-causing state; 86% (244/285) of those had a CADD score >15 (Supplementary File 3); 76% (214/285) of them had a CADD score >20. These included:

i. for genes labelled as monoallelic in PanelApp: 152 variants HGMD-listed variants with a CADD score >15 that were found to be present in heterozygous state in at least two individuals from the gnomAD control-only dataset.
ii. for genes labelled as biallelic in PanelApp: 48 variants HGMD-listed variants with a CADD score >15 that were found to be present in homozygous state in at least two individuals from the gnomAD control-only dataset.
iii. for genes labelled as both monoallelic and biallelic in PanelApp: 29 variants HGMD-listed variants with a CADD score >15 that were found to be present in homozygous state (in the case of variants reported to cause disease through a recessive mode of inheritance; n=6) or the heterozygous state (in the case of variants reported to cause disease through a dominant mode of inheritance; n=23) in at least two individuals from the gnomAD control-only dataset.
iv. for genes labelled as X-linked in PanelApp: 15 variants HGMD-listed variants with a CADD score >15 that were found to be present in either hemizygous or homozygous state in at least two individuals from the gnomAD control-only dataset.

### 1 in 6 genes implicated in inherited eye disorders is possibly associated with variable penetrance

Using the criteria outlined in the Methods section, we identified 56 IP/VE-associated genes (out of a total of 340 IED-implicated genes). Thirty-one of these were labelled as monoallelic, 11 as biallelic, 11 as both monoallelic and biallelic, and 3 as X-linked in PanelApp.

### Genes associated with variable penetrance exhibit variability in expression levels

Variability in gene expression at the population level was studied for IED-implicated genes. The GTEx v8 dataset was initially utilised: 75% (259/340) of genes were found to be adequately expressed in enough samples to allow the estimation of gene expression variability (the exact criteria are outlined in the Methods section); 43 of these were IP/VE-associated genes. The mean and median LCV scores were 57.1 and 60.9, respectively.

As retinal tissue is not included in the GTEx v8 dataset, the EyeGEx resource [18] was used to evaluate the subset of IED-implicated genes that has been linked to inherited retinal disorders (n=183). 88% (160/183) of genes were found to be adequately expressed in enough samples to allow estimation of gene expression variability; 23 of these were IP/VE-associated genes. The mean and median LCV scores were 62.7 and 71.8, respectively.

To test the hypothesis that variability in gene expression is associated with incomplete penetrance, we compared population variability in gene expression between IP/VE-associated genes and other IED-implicated genes (for which we did not find clear evidence of incomplete penetrance). First, all genes were considered and analysed based on the LCV scores in the entire GTEx dataset. IP/VE-associated genes with available scores (n=43) were significantly more variable than other genes (n=216) (median LCV, 83.7 vs. 48.85; p-value [Kolmogorov-Smirnov] = 0.0000015) when the median LCV of all tissues was used.

The same comparison was performed for clinically relevant and clinically actionable tissues. For example, when the median LCV of brain tissues was taken as the measure of variability, IP/VE-associated genes (n=35) were significantly more variable than other IED-implicated genes (n=184) (median LCV, 80.4 vs. 49.25; p-value [Kolmogorov-Smirnov] = 0.0000053). Similar results were obtained for whole blood (number of genes 15 vs. 101; median LCV 66.2 vs. 40.2; p-value [Mann-Whitney] = 0.018), sun-exposed skin (number of genes 22 vs. 156; median LCV 79.9 vs. 38.9; p-value [Mann-Whitney] = 0.0000083), non-sun-exposed skin (number of genes 23 vs. 152; median LCV 82 vs. 36.9; p-value [Kolmogorov-Smirnov] = 0.000017), and cultured fibroblasts (number of genes 23 vs. 152; median LCV 78.4 vs. 43.8; p-value [Mann-Whitney] = 0.00001).

In order to assess genes linked to inherited retinal disorders in more detail, we studied gene expression variability for this subset of IED-implicated genes using expression data from retinal tissue (from EyeGEx). IP/VE-associated retinal genes (n=23) were significantly more variable than other genes associated with inherited retinal disorders (n=137) (median LCV, 83.8 vs. 67.8; p-value [Kolmogorov-Smirnov] < 0.0028).

It is noteworthy that some genes commonly associated with variable penetrance in the literature were more variable than average at the mRNA level. Examples include *CYP1B1* (GTEx median LCV = 97.4), MYOC (GTEx median LCV = 98.5), *RP1L1* (EyeGEx LCV = 94.4), *BEST1* (EyeGEx LCV = 90.4), and *PRPF31* (EyeGEx LCV = 72).

All statistical comparisons in all the datasets and tissues assessed are shown in Table 1; the relevant plots are shown in Figure 2. We note that statistical significance was still observed after applying Bonferroni correction to the GTEx results, except for whole blood (although it should be noted that very few genes in the IP/VE group had LCV scores available in this tissue). Similar results were obtained when the analysis was repeated using the following variant-level cut-offs (instead of CADD score >15): CADD score >20, and CADD score >20 plus a minor allele frequency in gnomAD 2.1 of <0.01 (data not shown).

**Figure 2.**
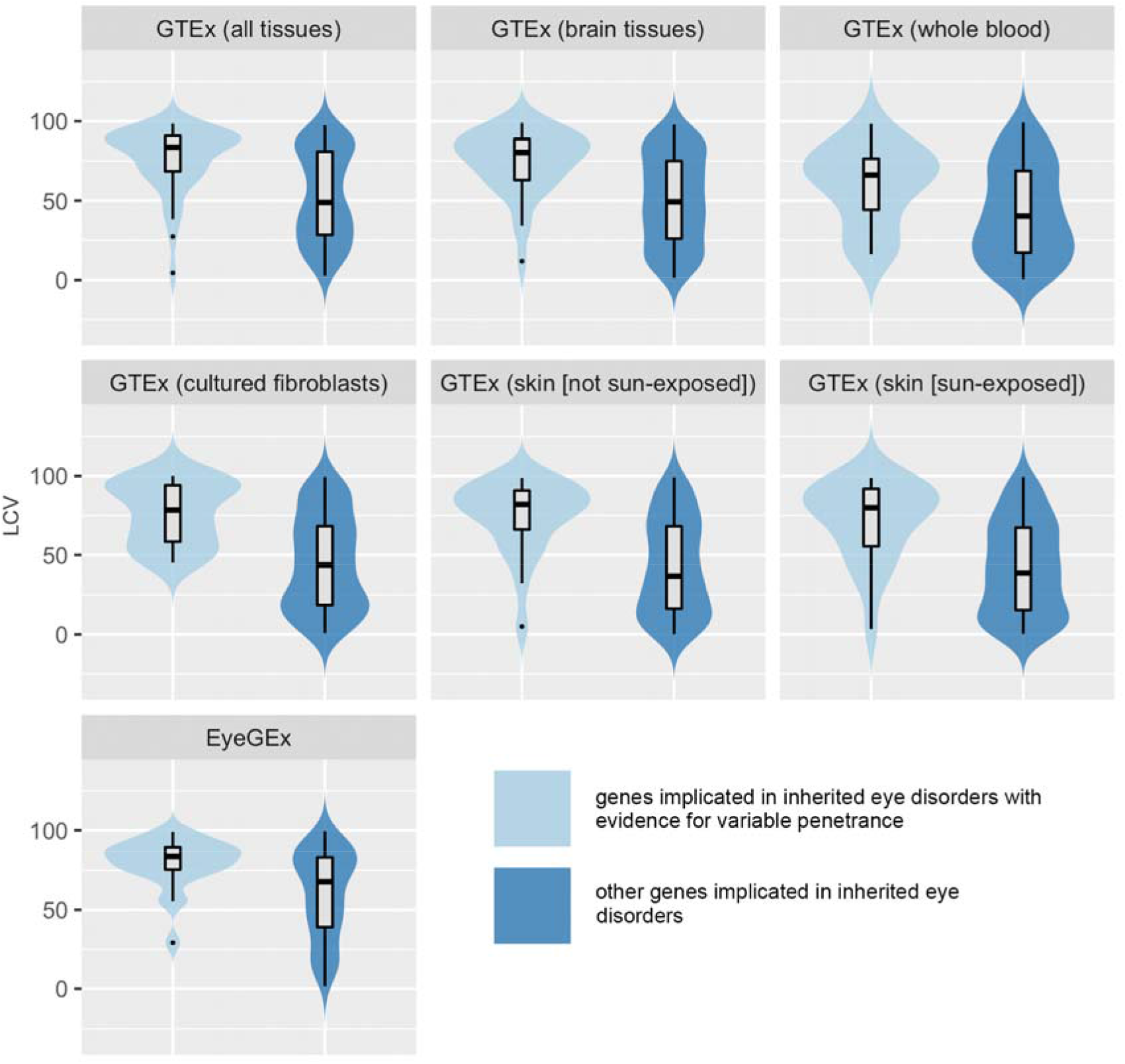
Violin plot showing comparisons of gene expression levels between inherited eye disease-implicated genes with evidence for variable penetrance and inherited eye disease-implicated genes for which no evidence of variable penetrance was found. The relevant p-values are presented in Table 1. LCV, local coefficient of variation; GTEx, Genotype-Tissue Expression Project version 8 dataset [9]; EyeGEx, Eye Genotype Expression (EyeGEx) dataset [18].

We then assessed whether the observed result was linked to differences in gene constraint. First, we looked for correlation between the LCV score and the missense o/e score in both the GTEx (all tissues and all genes) and EyeGEx (retinal tissue and genes implicated in inherited retinal disorders only) datasets; we found no significant correlation (Supplementary File 2). Second, we assessed the difference in various constraint metrics between IP/VE-associated genes and all other IED-implicated genes; the findings suggested that coding constraint is unlikely to be a significant confounding factor (Supplementary File 2).

To look for a possible “gradient effect”, we studied the relationship between the LCV score and the number of HGMD-listed variants that are present in the gnomAD control-only dataset for each gene; we found no obvious correlation (Supplementary File 2).

Gene ontology enrichment and interaction information for genes in the IP/VE group are provided in Supplementary file 2.

## Discussion

Advances in high-throughput sequencing technologies have enabled the creation of comprehensive catalogues of genetic variation (such as GTEx [9], gnomAD [17], and the UK Biobank [26]). Here, we used some of these population-scale datasets to study the hypothesis that pronounced phenotypic variation is associated with variability in gene expression; genes implicated in IED were used as a paradigm. We found that 1 in 6 IED-implicated genes have at least two possibly non-penetrant variants/alleles. We also found that the expression of these IP/VE-associated genes is significantly more variable than that of other IED-implicated genes (across tissues in general and in clinically relevant and actionable tissues in particular). These results highlight the potential value of incorporating both genomic variant information and gene expression level data in algorithms that aim to determine the precise risk of developing IED.

A number of studies have looked for overlap between databases cataloguing disease-associated variants (such as HGMD or ClinVar) and resources studying genetic variation at the population level (such as gnomAD or the UK Biobank). Although some of these studies evaluated the penetrance of disease-predisposing variants, the primary focus has typically been around questioning the validity of variant-disease assertions [27–32]. A study by Hanany and Sharon for example analysed the allele frequencies (in the full gnomAD dataset) of variants reported to cause autosomal dominant retinal disease (according to HGMD Public and RetNet [https://sph.uth.edu/retnet/]); the key finding was that 19% of genes and 10% of variants were spurious based on a carrier frequency threshold set by the authors [27]. Additionally, a recent study by Wright *et al.* performed a large-scale analysis of UK Biobank genotyping array data to assess the penetrance of ClinVar-reported variants in genes known to cause maturity-onset diabetes of the young (MODY) and severe developmental disorders; this led to a more refined penetrance estimate for some of the studied variants and to the refutation of a few previous disease associations [3]. Our work focused on a different group of disorders, IED, and had a number of differences in terms of study design. To reduce the potential impact of misannotation, we focused on [i] genes for which there is broad consensus for their involvement in disease [12] and [ii] variants that are considered to have strong evidence for pathogenicity (those marked as “DM” in HGMD) [13]. We chose to use HGMD (a pay-for-access resource which relies on curation of published literature) over ClinVar (a freely accessible public database which is based on submissions from researchers and clinical diagnostic laboratories) as, at the time of study design, the breadth of coverage of the former appeared to be greater [33]. Notably, a study that carefully curated >800,000 variants associated with hearing impairment found that variant classification discrepancies (compared to expert curation) were greater for ClinVar than for HGMD (14.5% and 5% respectively) [34]. Although these disease-associated variant resources will inevitably have some limitations, their ability to aggregate DNA alterations that are confidently linked to phenotypic abnormalities will continue to improve [35]. Combining them with population-scale variant datasets from carefully phenotyped cohorts (such as the UK Biobank) is expected to lead to the development of comprehensive catalogues of variants that are robustly associated with variable penetrance.

Previous studies have used allele specific expression (rather than total expression) to understand how variability in gene expression impacts variant penetrance [6,10]. Allele-specific expression analysis allows imbalances between the expression of the two relevant alleles to be assessed, thus identifying large-effect *cis*-acting regulatory elements. However, there are examples of regulatory variants that modulate the penetrance of variants causing IED in *trans* [1,7]. We attempted to account for overall variation by assessing the total mRNA count for each gene and by calculating variability in large populations of individuals that are presumably not affected by Mendelian disorders (GTEx [9] and EyeGEx [18]). Future studies could build on these results by developing a comprehensive ranking of IED-implicated genes based on their variation in gene expression as the result of *trans-* and *cis*-acting elements.

This study has a number of notable limitations. First, penetrance and expressivity characterise the relationship between a genotype and a phenotype, while gene expression is a characteristic of a gene; its variation therefore acts only as a proxy for variable expressivity in the case of individual variants. Second, we did not consider the probability of a gene being associated with gain-of-function or loss-of-function (or dosage sensitivity) effects; future studies incorporating data on the pathophysiology underlying each molecule are expected to provide further insights. Third, gene expression for each given IED-implicated gene needs to be assessed in the relevant disease target tissue [36]. It is however not always possible to obtain comprehensive data for the desired tissue type. We aimed to limit the impact of this by assessing the median value across multiple tissues and by confirming the findings for retinal disease-associated genes in a dedicated expression dataset from donor retinas. Future work will utilize increasingly powerful RNA-sequencing-based transcriptome resources of healthy human eye tissues such as eyeIntegration [37]. Finally, it is worth highlighting that association does not necessarily suggest causation: our findings do not demonstrate a mechanistic link between gene expression variability and variable penetrance although they provide an additional line of evidence to support this hypothesis.

In conclusion, we determined a subset of genes implicated in IED that are associated with a degree of variable penetrance. Many genes in this subset have highly variable gene expression at the population level. It can be speculated that this finding reflects the involvement of *cis* or *trans*-acting genetic regulatory modifiers. Further studies utilising large-scale genomic and transcriptomic datasets are expected to provide important insights into the precise nature of these regulatory modifier mechanisms and to highlight in-human validated targets for novel treatments.

## Acknowledgments

We acknowledge the following sources of support and funding: Christopher Green and the NIHR Clinical Lecturer Programme (CL-2017-06-001, PIS). We thank the two anonymous reviewers for their constructive comments, which helped us to improve this manuscript.

