## Supplementary file 2 for "Variability in gene expression is associated with incomplete penetrance in inherited eye disorders"

Variability in gene expression is associated with incomplete penetrance in inherited eye disorders

David J. Green^1^, Shalaw R. Sallah^1^, Jamie M. Ellingford^1,2^, Simon C. Lovell^1^, Panagiotis I. Sergouniotis^1,2,3^

^1^ Division of Evolution and Genomic Sciences, School of Biological Sciences, Faculty of Biology, Medicines and Health, University of Manchester M13 9PT, UK.

^2^ Manchester Centre for Genomic Medicine, St Mary's Hospital, Manchester University NHS Foundation Trust, Manchester M13 9WL, UK.

^3^ Manchester Royal Eye Hospital, Manchester University NHS Foundation Trust, Manchester M13 9WL, UK.

**Figure S1**. Scatter plot showing the relationship between the Genotype-Tissue Expression Project (GTEx) and the Eye Genotype Expression (EyeGEx) local coefficient of variation (LCV) scores in genes that are associated with inherited eye disorders and are adequately expressed in both datasets (*r* = 0.441).

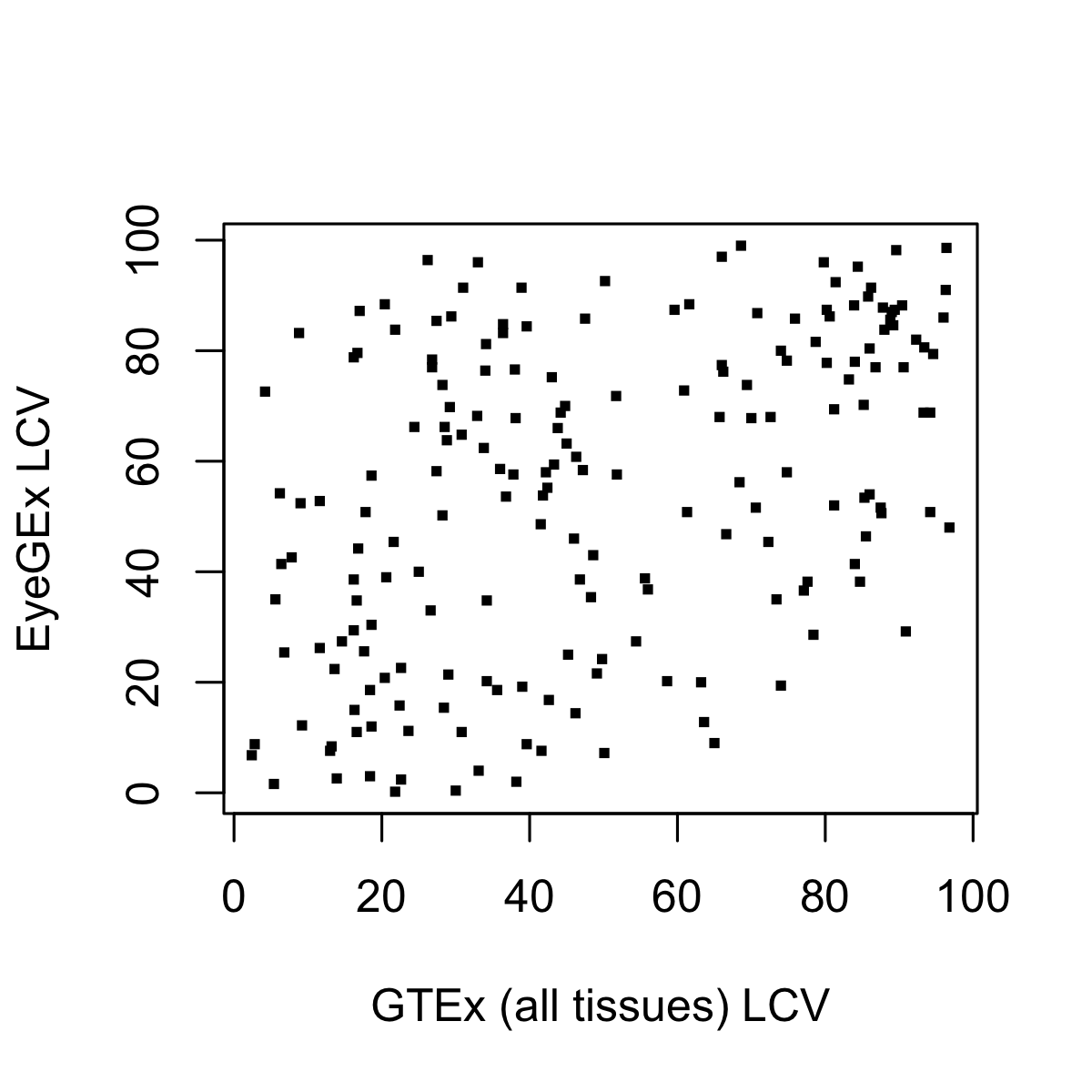

**Figure S2.** Scatter plot showing the relationship between the Genome Aggregation Database (gnomAD) missense observed/expected (o/e) score and the Genotype-Tissue Expression Project (GTEx) local coefficient of variation (LCV) score (*r* = 0.106).

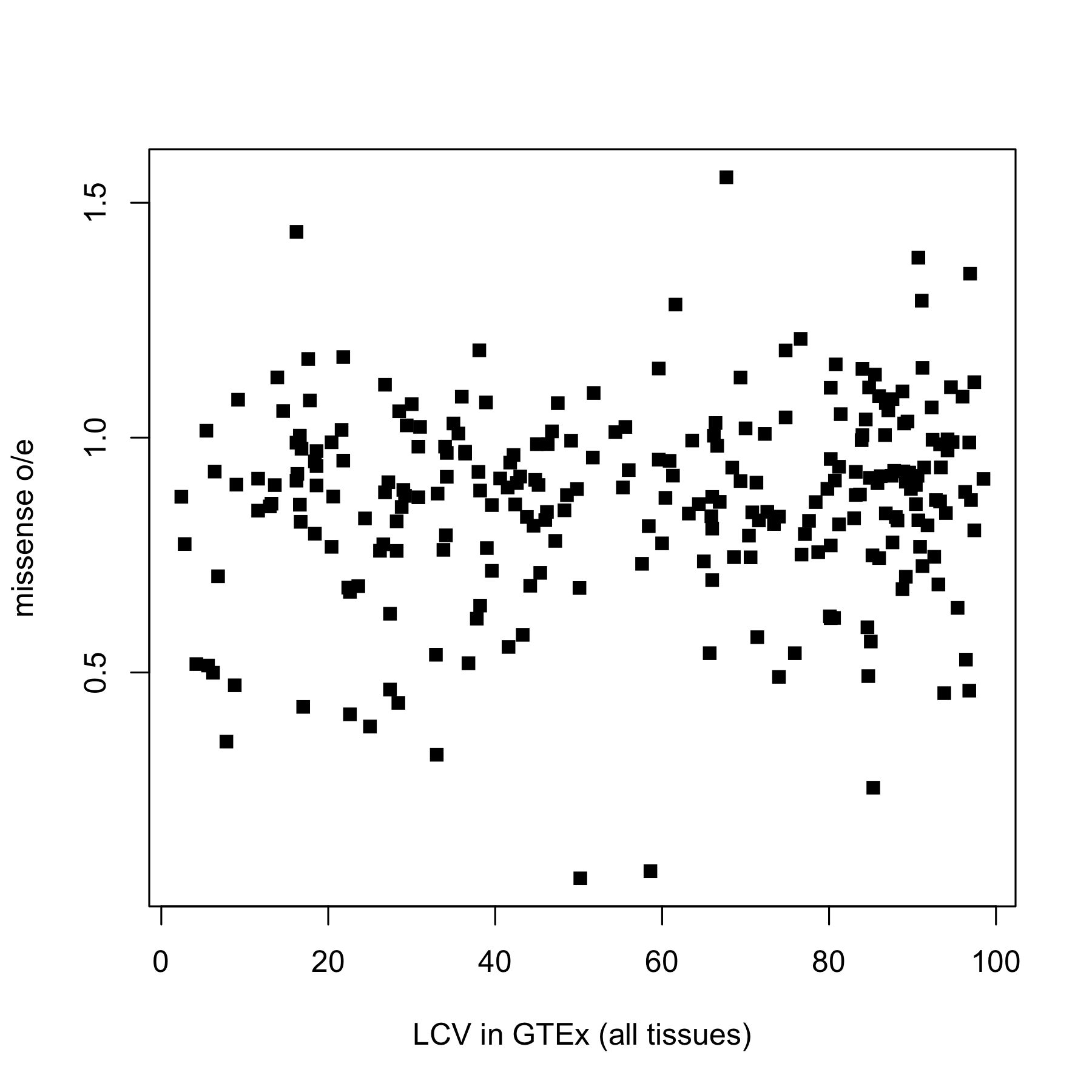

**Figure S3.** Scatter plot showing the relationship between the Genome Aggregation Database (gnomAD) missense observed/expected (o/e) score and the Eye Genotype Expression (EyeGEx) local coefficient of variation (LCV) score (*r* = 0.175).

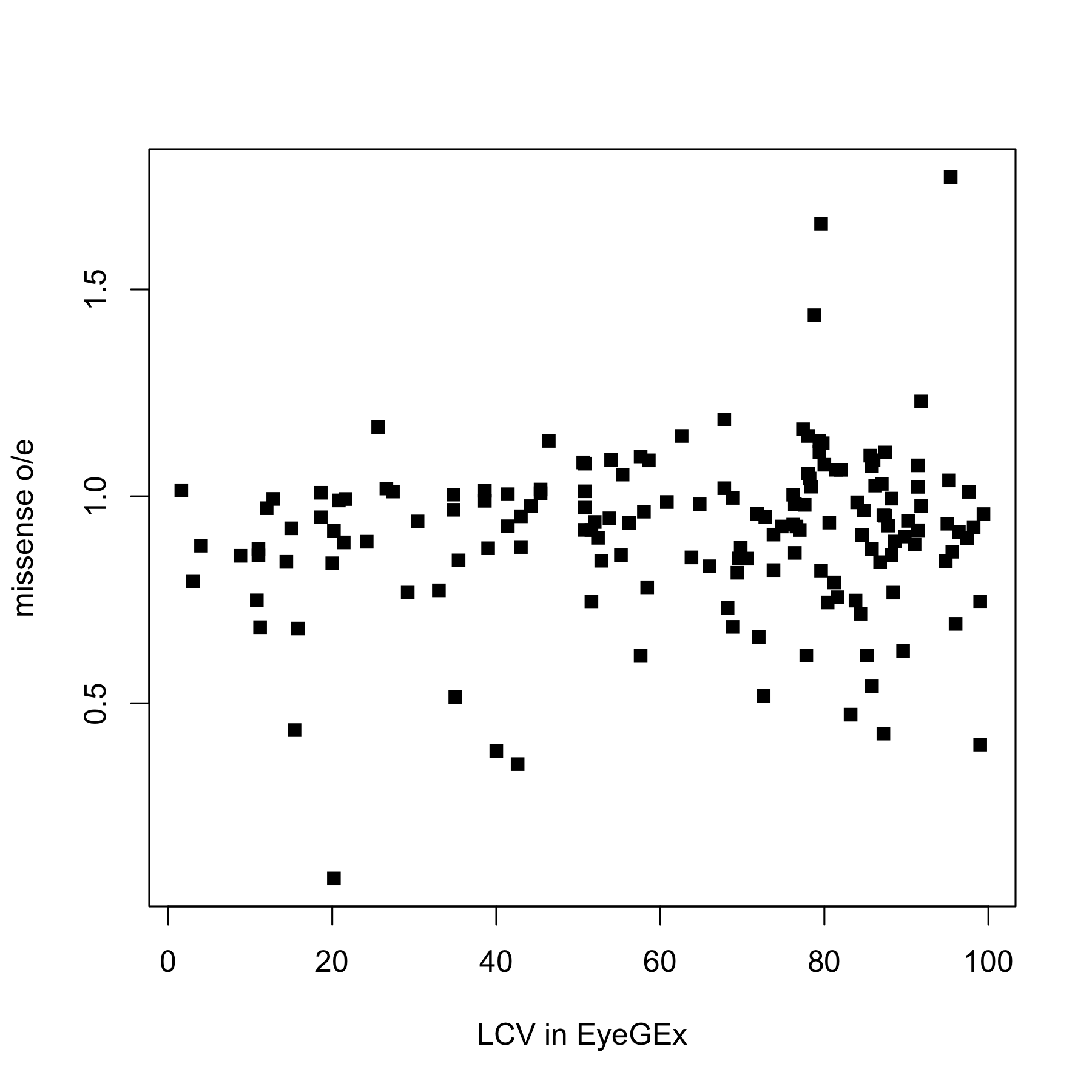

| **Table S1.** Comparison of gene constraint metrics between genes that are possibly associated with variable penetrance and genes for which no evidence of variable penetrance was detected. | | | | |
| --- | --- | --- | --- | --- |
| **metric** (as per gnomAD v2.1.1) | **mode of inheritance**  (as per PanelApp) | **IP/VE IED genes** | **other IED genes** | **p-value** |
| missense o/e | all | 0.891 | 0.906 | 0.1 |
| missense Z | all | 0.951 | 0.569 | 0.044 |
| LOF o/e | all | 0.420 | 0.547 | 0.046 |
| LOEUF | all | 0.658 | 0.858 | 0.008 |
| missense o/e | monoallelic | 0.811 | 0.711 | 0.042 |
| missense Z | monoallelic | 1.489 | 1.631 | 0.262 |
| LOF o/e | monoallelic | 0.190 | 0.189 | 0.446 |
| LOEUF | monoallelic | 0.434 | 0.432 | 0.342 |
| missense o/e | biallelic | 0.996 | 0.948 | 0.139 |
| missense Z | biallelic | 0.073 | 0.341 | 0.228 |
| LOF o/e | biallelic | 0.673 | 0.600 | 0.153 |
| LOEUF | biallelic | 0.848 | 0.908 | 0.324 |
| missense o/e | both monoallelic and biallelic | 0.941 | 0.900 | 0.069 |
| missense Z | both monoallelic and biallelic | 0.359 | 0.591 | 0.121 |
| LOF o/e | both monoallelic and biallelic | 0.583 | 0.513 | 0.119 |
| LOEUF | both monoallelic and biallelic | 1.013 | 0.894 | 0.178 |
| missense o/e | X-linked | 0.748 | 0.793 | 0.376 |
| missense Z | X-linked | 1.879 | 1.082 | 0.563 |
| LOF o/e | X-linked | 0.181 | 0.087 | 0.684 |
| LOEUF | X-linked | 0.448 | 0.314 | 0.244 |
| gnomAD, the Genome Aggregation Database; IED, inherited eye disease; IP/VE IED genes, IED-implicated genes that are enriched for variants with possible variable penetrance; other IED genes, IED-implicated genes that are not known to be enriched for variants with possible variable penetrance; missense o/e, missense observe/expected score; missense Z, missense z-score; LOF o/e, loss-of-function observed/expected score; LOEUF, loss-of-function observed/expected upper bound fraction.  For missense o/e, LOF o/e and LOEUF a lower score suggests a high constraint or intolerance to variation. For missense Z, a higher score suggests a high constraint or intolerance to variation.  The p-values shown are before adjusting for multiple comparisons. | | | | |

**Figure S4.** Scatter plots showing the relationship between the local coefficient of variation (LCV) in GTEx (median of all tissues) and the number of per-gene HGMD-listed variants that are present in the gnomAD control-only dataset*.* Only data on a set of inherited eye disease-implicated genes with evidence for variable penetrance are shown. Variants in overlap, number of HGMD-listed variants with CADD >15 that are present in the gnomAD control-only dataset; variants in overlap/starting variants, number of HGMD-listed variants with CADD >15 that are present in the gnomAD control-only dataset corrected for the overall number of HGMD-listed variants in the relevant gene; variants in overlap/coding length, number of HGMD-listed variants with CADD >15 that are present in the gnomAD control-only dataset corrected for canonical protein coding length (CDS) for the relevant HGMD transcript. HGMD, Human Gene Mutation Database.

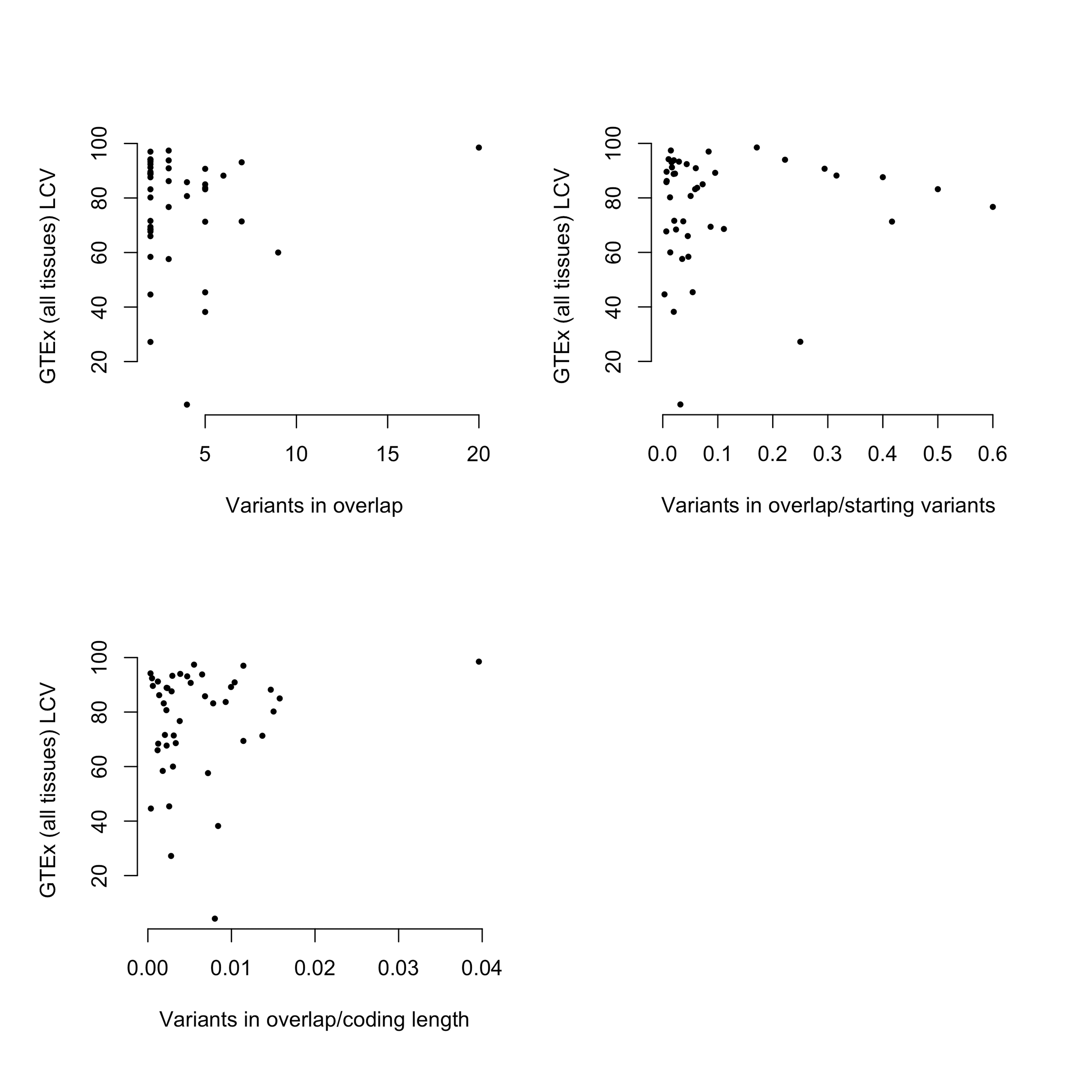

**Figure S5**. Scatter plots showing the relationship between the local coefficient of variation (LCV) in EyeGEx and the number of per-gene HGMD-listed variants that are present in the gnomAD control-only dataset*.* Only data on a set of inherited retinal disease-implicated genes with evidence for variable penetrance are shown. Variants in overlap, number of HGMD-listed variants with CADD >15 that are present in the gnomAD control-only dataset; variants in overlap/starting variants, number of HGMD-listed variants with CADD >15 that are present in the gnomAD control-only dataset corrected for the overall number of HGMD-listed variants in the relevant gene; variants in overlap/coding length, number of HGMD-listed variants with CADD >15 that are present in the gnomAD control-only dataset corrected for canonical protein coding length (CDS) for the relevant HGMD transcript.

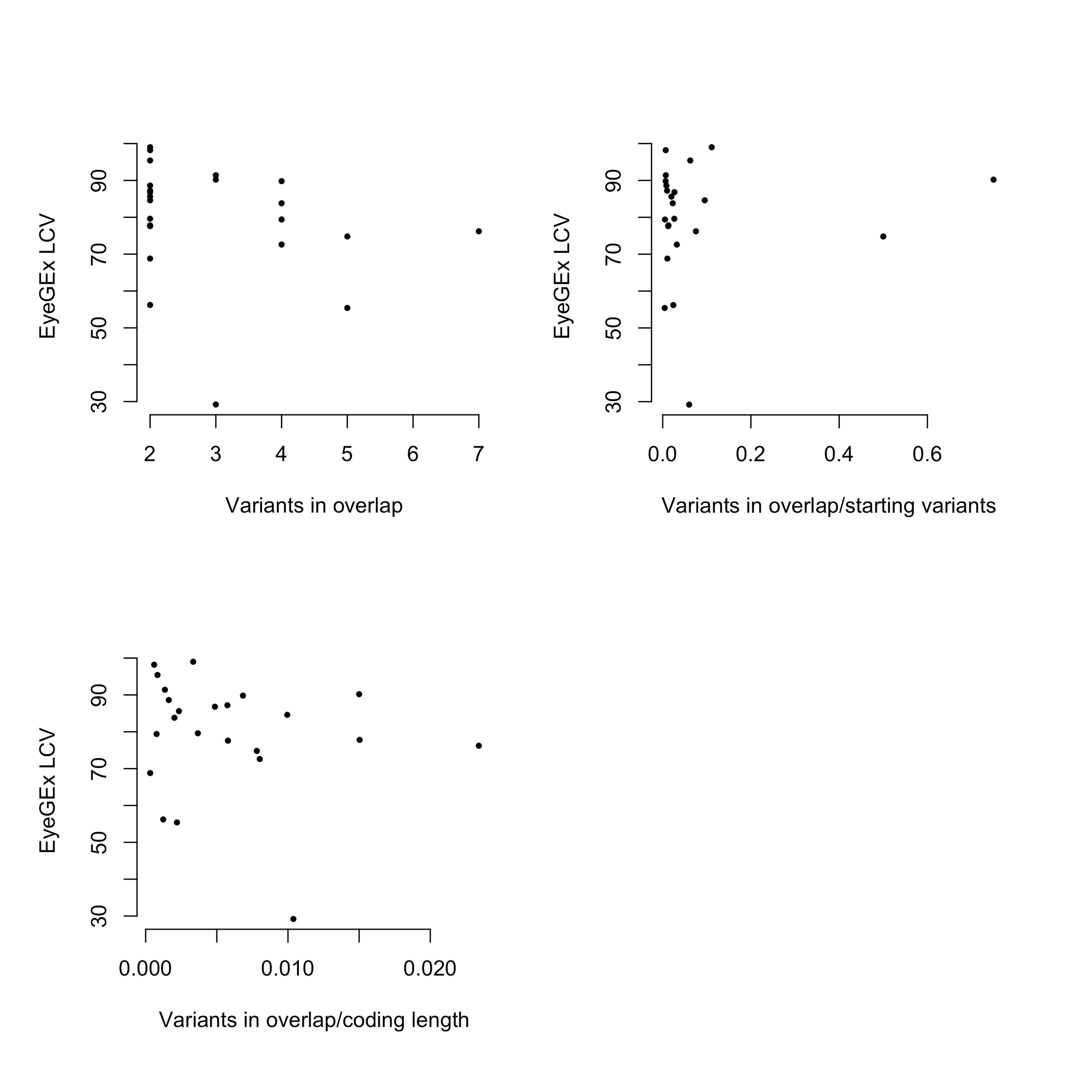

| **Table S2.** Gene ontology (GO) term enrichment performed with STRING v11 [37] for the most variable (in terms of location coefficient of variation) genes associated with variable penetrance (top 10 most variable genes in the Genotype-Tissue Expression Project and top 10 most variable genes in the Eye Genotype Expression datasets [genes implicated in inherited retinal disease only]); the top 20 most significantly enriched GO terms are shown. | | | |
| --- | --- | --- | --- |
| **GO term ID** | **GO term description** | **observed gene count** | **false discovery rate** |
| 0007601 | visual perception | 12 | 5.24x10^-16^ |
| 0007423 | sensory organ development | 12 | 5.94x10^-12^ |
| 0001654 | eye development | 10 | 1.51x10^-10^ |
| 0043010 | camera-type eye development | 9 | 1.19x10^-9^ |
| 0009583 | detection of light stimulus | 6 | 5.38x10^-9^ |
| 0003008 | system process | 14 | 1.22x10^-8^ |
| 0007602 | phototransduction | 5 | 1.21x10^-7^ |
| 0048513 | animal organ development | 15 | 2.84x10^-7^ |
| 0060041 | retina development in camera-type eye | 6 | 3.93x10^-7^ |
| 0022400 | regulation of rhodopsin mediated signalling pathway | 4 | 1.27x10^-6^ |
| 0009314 | response to radiation | 7 | 1.10x10^-5^ |
| 0032501 | multicellular organismal process | 18 | 1.60x10^-5^ |
| 0050896 | response to stimulus | 18 | 3.00x10^-4^ |
| 0045494 | photoreceptor cell maintenance | 3 | 4.40x10^-4^ |
| 0009653 | anatomical structure morphogenesis | 10 | 4.50x10^-4^ |
| 0001525 | angiogenesis | 5 | 5.40x10^-4^ |
| 0009584 | detection of visible light | 3 | 6.70x10^-4^ |
| 0042461 | photoreceptor cell development | 3 | 6.70x10^-4^ |
| 0061304 | retinal blood vessel morphogenesis | 2 | 6.70x10^-4^ |
| 0001894 | tissue homeostasis | 4 | 7.30x10^-4^ |

**Figure S6.** Interaction networks for the 10 most variable (in terms of local coefficient of variation) genes associated with variable penetrance in (A) the Genotype-Tissue Expression Project and (B) the Eye Genotype Expression (genes implicated in inherited retinal disease only) datasets. Only high-confidence interactions are shown (minimum interaction score = 0.7); networks were built using the STRING v11 web server [37].

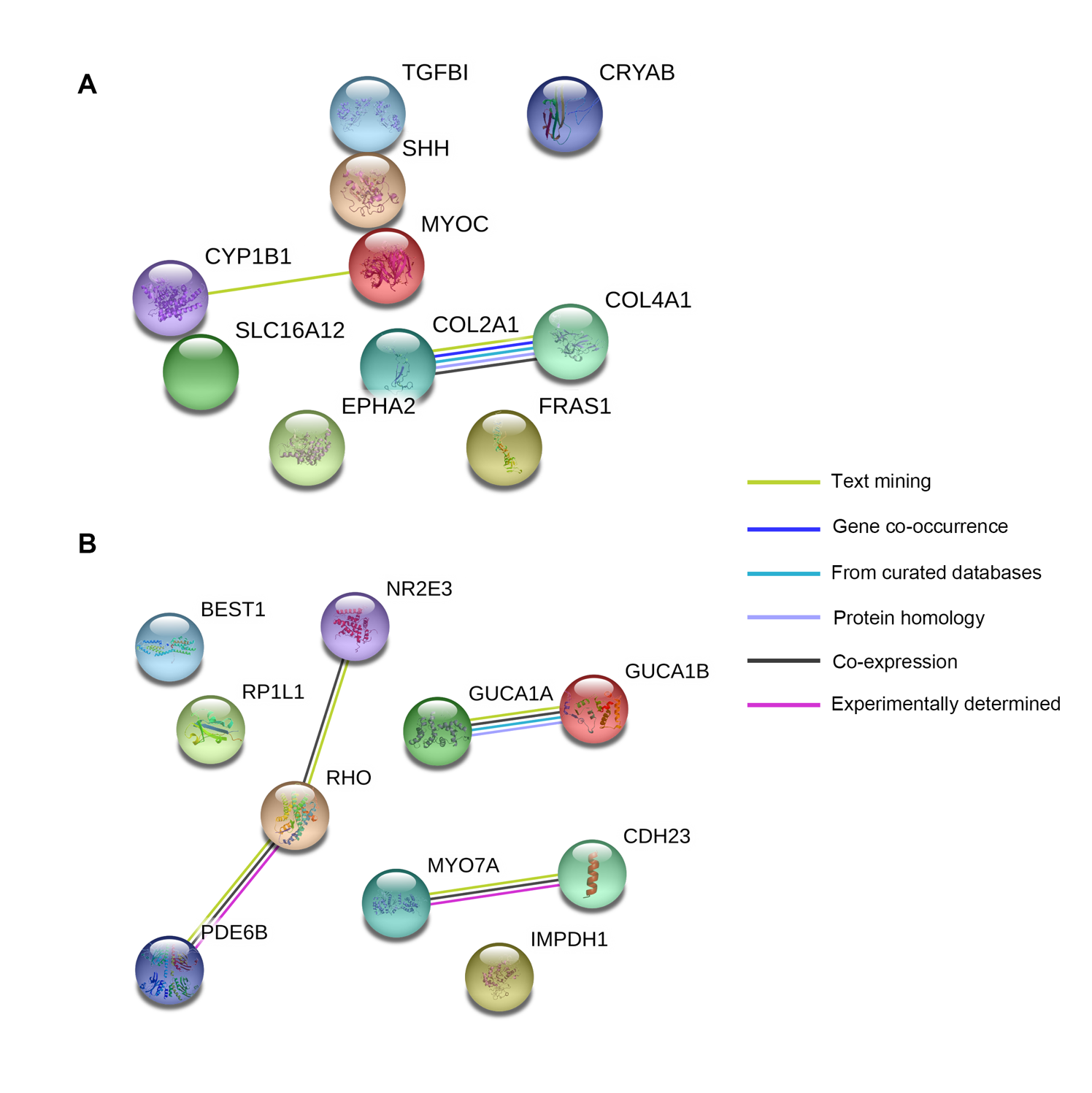
